# Galectin-3 recruitment at the *Mycobacterium tuberculosis*-containing phagosome is critical in macrophage but dispensable in epithelial cells

**DOI:** 10.64898/2026.06.04.730029

**Authors:** Yoël Dagan, Nathalie Deboosere, Eugénie Boulagnon, Aurélie Burette, Arnaud Machelart, Olivier Molendi-Coste, Sophie Desnoulez, Elisabeth Werkmeister, Roxane Simeone, Alexandre Grassart, Priscille Brodin

## Abstract

*Mycobacterium tuberculosis* (*Mtb*) virulence relies in part on its ability to induce phagosomal membrane rupture, enabling bacterial access to the host cell cytosol. This process is largely mediated by the ESX-1 secretion system, which is present in *Mtb* but absent from the vaccine strain BCG. Galectin-3 (Gal3), a β-galactoside–binding lectin, is recruited to damaged endomembranes and functions as a cytosolic sensor of membrane disruption. However, the kinetics and quantitative features of Gal3 recruitment to *Mtb*-containing vacuoles have remained poorly characterized.

Here, we performed a longitudinal quantitative imaging study of Gal3 recruitment in human macrophages over a five-day infection period. Gal3 was recruited to mycobacteria-containing vacuoles shortly after infection with both live and heat-killed *Mtb*. Differences between the two conditions emerged from day 1 post-infection and persisted until macrophage death. A similar kinetic profile was observed with recombinant BCG::ESX-1, whereas parental BCG failed to induce Gal3 recruitment, confirming the requirement for ESX-1–dependent membrane damage.

Cytoplasmic Gal3 levels were higher in bystander macrophages than in infected cells and were comparable to non-infected controls, suggesting that diffuse cytoplasmic Gal3 is associated with cells lacking intracellular mycobacteria. Functional studies revealed that Gal3 silencing enhanced long-term intracellular *Mtb* replication, demonstrating a role for Gal3 in restricting bacterial growth.

Importantly, Gal3 recruitment was not observed in alveolar epithelial cells. This cell-type specificity was confirmed in a microfluidic alveolus-on-chip model. Together, these findings identify sustained Gal3 recruitment to the mycobacteria-containing vacuole as a robust quantitative marker of ESX-1–dependent phagosomal rupture and *Mtb* virulence.

## Introduction

Tuberculosis (TB), caused by *Mycobacterium tuberculosis* (*Mtb*), remains a major global health challenge. In 2024, it was the leading cause of death from a single infectious agent, accounting for approximately 1.23 million deaths worldwide, underscoring its continued public health burden [1]. Despite antibiotic therapeutic strategies, the emergence and spread of drug-resistant *Mtb* strains have increasingly limited the effectiveness of these conventional treatments. Consequently, a better understanding of the fundamental mechanisms in *Mtb* infection for the development of new therapeutic strategies and new regimen remains a critical priority for global tuberculosis control.

Within the lung alveoli, *Mtb* exhibits a strong tropism for alveolar macrophages, which represent its primary intracellular niche for replication and long-term persistence. Nevertheless, other alveolar cell types, including type I and type II epithelial cells, were found to be also infected [2–6]. The functional consequences of this cellular host heterogeneity for bacterial dissemination and raises the question about the influence of the complex multicellular alveolar microenvironment on the outcome during early events of infection.

A key step in *Mtb* pathogenesis is its confinement and escape from a specific intracellular structure of the host cell known as the phagosomal vacuole. Over the last two decades, several studies based on electronic microscopy analysis and CCF4 FRET photonic microscopy, monitored phagosomal membrane rupture during the course of *Mtb* infection and showed that this phenomenon was dependent of the mycobacterial lipid PDIM and ESX-1 secretion system [7–9]. The dependency of ESX-1 has been also confirmed by electron microscopy in both *Mycobacterium marinum* and *M. tuberculosis* and importantly phagosomal rupture has been also detected *in vivo*, supporting the biological relevance of this process in *Mtb* infection [10, 11]. Taken together, these findings established that phagosomal escape associated by membrane damage plays a pivotal role of mycobacterial virulence.

Beyond *M. tuberculosis*, elucidating the mechanisms underlying vacuolar escape is of broad importance, as this conserved step is essential for the infectivity of many bacterial pathogens, including *Shigella*, *Listeria*, and *Salmonella* [12, 13]. Interestingly, the cytosolic β-galactoside–binding lectin Galectin-3 (Gal3) has emerged as potential common sensors for detecting endomembrane damage. Briefly, under physiological conditions, luminal glycoconjugates are confined within intact vacuoles but recruited to disrupted vacuoles upon bacterial escape, reflecting cytosolic exposure of these glycan. Gal3 has primarily been investigated during the early stages of *Mtb* infection, particularly within the first minutes to several hours post-infection [14–17]. In these studies, Gal3-positive puncta colocalizing with *Mtb* were observed in approximately 15% of infected phagocytes within the cellular monolayer.

The use of Gal3 as a marker of vacuolar rupture has facilitated the identification of host factors involved in this process, including *Mtb* infection, such as VPS18 [15]. Despite growing interest in host determinants of *Mtb* infection, relatively few factors have been mechanistically linked to phagosome rupture, and their temporal dynamics, cell-type specificity, and functional hierarchy remain incompletely understood [15, 18, 19]. To address these gaps, experimental systems are required to capture both the kinetics of intracellular events and the complexity of the pulmonary microenvironment.

Beyond its use as a marker for endomembrane damages, several evidence indicates that Gal3 could be directly involved in the host cell response, including the activation of the ESCRT pathway and the promotion of membrane repair of damaged lysosomes [20]. But most notably, it has been shown to participate directly in host defense pathways, including inflammasome activation and infection-induced programmed cell death, suggesting that it may function not only as a endomembrane damage marker but also as an effector in antimicrobial responses [13, 21]. However, the precise role of Gal3 during *Mtb* infection, especially in relation to phagosomal rupture events, remains to be better understood.

By integrating high-content imaging, genetic perturbation and organotypic modeling, our study aims to characterize the dynamics and functional significance of Gal3 expression in *Mtb* infection. Briefly, two complementary strategies were implemented. First, we performed longitudinal analyses of phagosomal damage and host responses across multiple time points using siRNA-based perturbation in the widely used macrophage cell line THP-1. Second, we used a recently human Alveolus-on-Chip (AoC) model developed in our lab [22], to investigate phagosomal rupture within a multicellular architecture that more closely resembles to the *in vivo* alveolar barrier.

## Results

### Galectin-3 spots sustainably colocalizes with *Mtb* for several days in macrophages after infection

We analyzed Gal3 dynamics over a five-day infection course in THP-1 differentiated human macrophages with a virulent red fluorescent strain of *Mtb* (H37Rv) followed by immunostaining with an anti-Galectin 3 (Gal3) antibody and nuclear labeling with DAPI (Figure 1A). At the cellular level, Gal3 displayed a broad cytoplasmic distribution as well as discrete punctate structures, then referred here as Gal3 “spots” defined by signal intensities at least two-fold higher than the surrounding cytoplasm (Figure 1B). These Gal3 spots have previously been associated with membrane damage or, when colocalizing with *Mtb*, with phagosomal rupture [14]. Colocalization was not observed for heat-killed (HK H37Rv) used as control (Figure 1B). We therefore quantified the proportion of bacteria colocalizing with Gal3 spots, here defined as Gal3-positive H37Rv (Figure 1C). Approximately 20% of both live and HK H37Rv colocalized with Gal3 after 3 hours post-infection, whereas this proportion decreased and stabilized at lower levels over subsequent days. From one day post-infection onward, live bacteria exhibited significantly higher Gal3 colocalization compared with HK controls. These findings suggest that early Gal3 recruitment is likely driven by phagocytic uptake and associated membrane remodeling, while sustained recruitment reflects active bacterial processes, indicating that Gal3 engagement extends beyond the initial stages of infection in this model.

**Fig 1.**
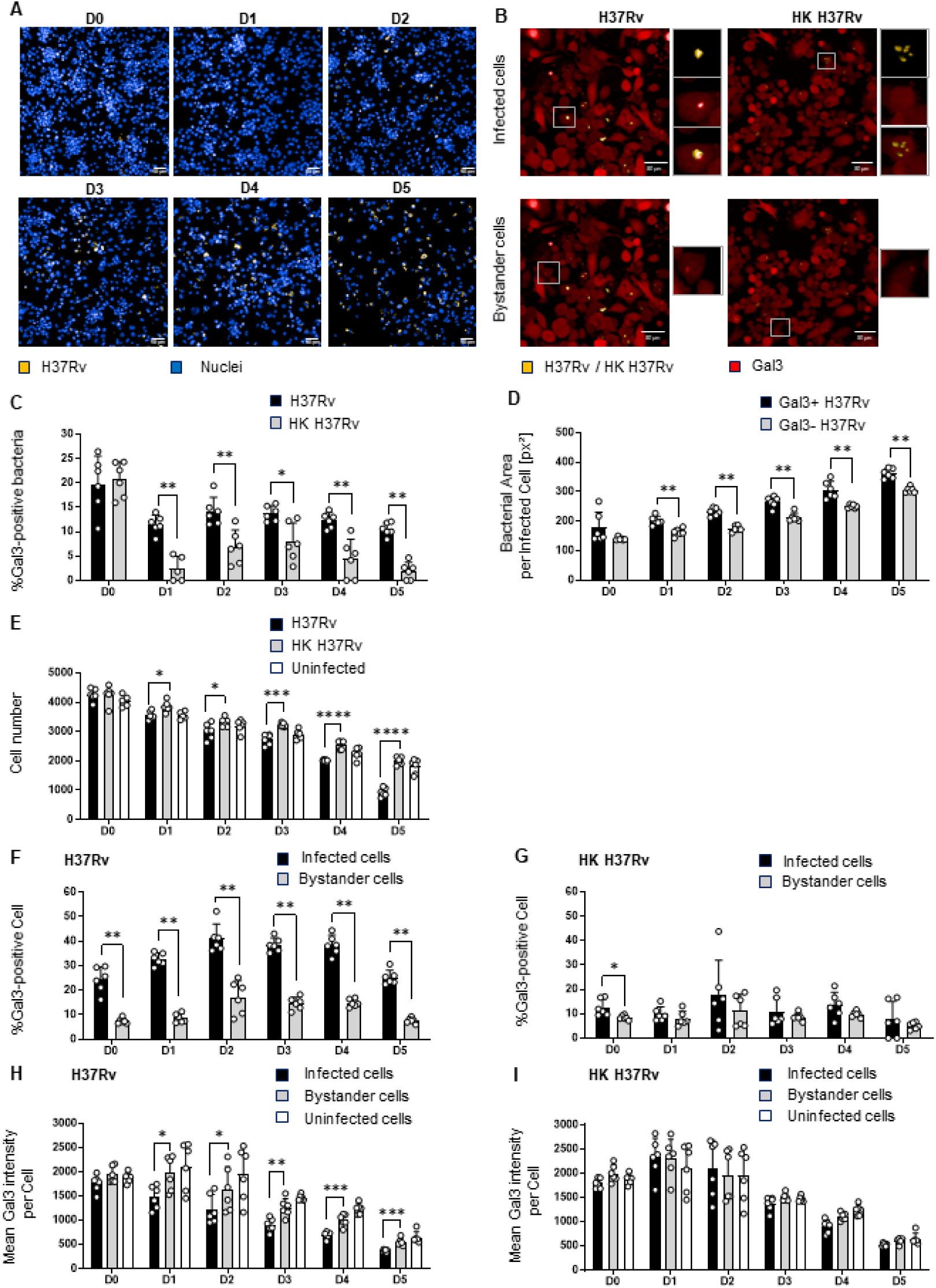
Galectin-3 recruitment to intracellular bacteria and associated cellular phenotypes during infection. THP-1 cells were grown in 384-well plates, infected with Mtb H37Rv pMRF1 (MOI = 1) or heat-killed (HK) H37Rv pMRF1 for 5 days and imaged by automated confocal microscopy. (A) Representative images of THP-1 (nucleus; blue) infected with H37Rv pMRF1 (yellow) over 5 days. Scale bar = 50 μm (20X). (B) Representative images of THP-1 infected with live and HK H37Rv pMRF1 (MOI = 1; yellow) and labelled by immunofluorescence for Galectin-3 (Gal3; red) at day 3 post-infection. (Left) The focused image highlights a live bacterium colocalized with a Gal3 spot, indicating phagosomal membrane damage, while (right) no colocalization of HK bacterium with Gal3 spot is shown. The magnified inset of bystander cells under conditions of infection with both live and HK bacteria show Gal3 spots, suggesting cellular damage. Scale bar = 50 μm (20X). (C, D, E, F, G) Quantitative image analysis of infected and bystander HPAEpiC cells infected with live and HK H37Rv pMRF1 and in non-infected conditions over 5 days. (C) Percentage of bacteria colocalizing with Gal3 spots. Live bacteria display significantly higher rate of colocalization with Gal3 than HK bacteria from day 1 post-infection (Mann-Whitney, P≤0.05). (D) Intracellular bacterial area (pixel²) per infected cells containing at least one bacterium that is positive for Gal3 (Gal3+, rupture) compared to cells infected with bacteria not colocalized with Gal3 spot (Gal3-, non-rupture). The cells exhibiting colocalization with Gal3 show significantly greater bacterial area from 1 day after infection (Wilcoxon test, P≤0.05). (E) Total cell number. No significant difference is detected on day 0, while cell numbers obtained for live-infection conditions and HK controls are significantly different from day 1 post-infection (One-way Anova, P≤0.05). (F) Percentage of infected and bystander Gal3-positive THP-1 cells for infection with live bacteria. Infected cells show significant higher rate of Gal3-positive cells than bystander cells (Paired t-test, P≤0.05). (G) Percentage of infected and bystander Gal3-positive THP-1 cells for infection with HK bacteria. Infected cells exhibit a significantly higher proportion of Gal3-positive cells up to the first day following infection, whereas no significant difference is observed from the day 2 (Paired t-test, P≤0.05). (H) Mean cellular Gal3 fluorescence intensity. Infected cells exhibit significantly reduced Gal3 intensity relative to bystanders from day 1 (Paired t-test, P≤0.05), whereas bystander and uninfected cells do not differ (Unpaired t-test). (I) Mean cellular Gal3 fluorescence intensity over time in infected and bystander cells obtained for infection with HK H37Rv pMRF1, and in uninfected controls. No significant differences (paired and unpaired t-test, P≤0.05).

Importantly, Gal3 recruitment to live bacteria was associated with a significantly increased bacterial area per cell, based on the measure of the number of pixels of detected bacteria in each infected cell (Figure 1D). This supports a link between Gal3 accumulation, phagosomal membrane rupture, and intracellular bacterial replication that has been assessed observing the bacterial area per infected cell through microscopy [23]. This correlation became apparent from day 2 post-infection, consistent with the known slow replication kinetics characteristic of *Mtb* (Figure 1D). Across five days of infection, a progressive decline in cell numbers was observed in all conditions, including HK control (Figures 1E). As differentiated THP-1 macrophages do not proliferate, this decrease likely reflects both the cellular infection and the intrinsic cell apoptosis. Notably, infection with live H37Rv resulted in significantly increased cell death, detectable as early as one day post-infection, with the most pronounced loss occurring between days 4 and 5. Together, these results highlight a stable temporal relationship between Gal3 recruitment, phagosomal damage, and bacterial growth during macrophage infection.

### H37Rv induces alterations of Galectin-3 in bystander cells

In addition to assessing Gal3 recruitment at bacterial colocalization sites, we analyzed its global distribution in infected and non-infected cells independently of direct bacterial association. Cells exhibiting Gal3 spots were defined as Gal3-positive (Figure 1B). Following infection with live *Mtb*, a significantly higher proportion of Gal3-positive cells was observed among infected cells compared with bystander cells (Figure 1F). In contrast, infection with heat-killed (HK) bacteria did not result in any significant difference between infected and bystander populations (Figure 1G). These findings are consistent with the notion that Gal3 spot formation is associated with host cell damage induced by live bacteria and is likely partly driven by secreted virulence factors, such as ESAT-6, rather than by bacterial membrane components alone, such as PDIM.

We next quantified the total intracellular Gal3 signal intensity across conditions. Infected cells showed a significant reduction in overall Gal3 expression compared to bystander cells starting one day after infection, in contrast with what was observed in cells exposed to HK bacteria (Figure 1H-I). Given the substantial Gal3 recruitment to bacteria at early time points, the decrease cannot be attributed to a redistribution of Gal3 within the cell. This difference was observed over the course of five days post-infection, suggesting an infection-dependent modulation of Gal3 abundance. The progressive reduction in Gal3 levels observed in each condition over time may indicate that cell death also reduces Gal3 expression.

### Similar phenotype observed using a derivative BCG::ESX-1 *Mycobacterium marinum* strain by comparison with a BCG strain

To further investigate the involvement of the type VII secretion system ESX-1 in the observed phenotype, we performed comparable analysis using the non-virulent mycobacterial strains BCG and BCG::ESX-1 *Mycobacterium marinum* (*Mmar*). The BCG::ESX-1*^Mmar^* strain possesses a functional ESX-1 secretion system, obtained by introducing the *Mycobacterium marinum* RD1 locus. This locus enables expression of the ESX-1 type VII secretion system and has been shown to confer phagosomal rupture proficiency comparable to that of *Mtb*, while exhibiting reduced virulence [24, 25]. Over a six-day infection period, we observed a pattern of Gal3 recruitment comparable to that seen with live H37Rv versus HK H37Rv, when comparing BCG::ESX-1*^Mmar^*and BCG respectively (Figures 2A-C). A significantly higher proportion of Gal3 spot colocalization was also detected in cells infected with BCG::ESX-1*^Mmar^* compared with BCG-infected cells, starting 2 days post-infection. No significant difference was observed at the time of infection or one day later, when approximately 20% of bacteria in both conditions colocalized with Gal3, further supporting the notion that early Gal3 recruitment is largely driven by phagocytic uptake and associated membrane remodeling. When looking at the number of cells over the course of the infection, no significant difference was observed between the different conditions (Figure 2C).

**Fig 2.**
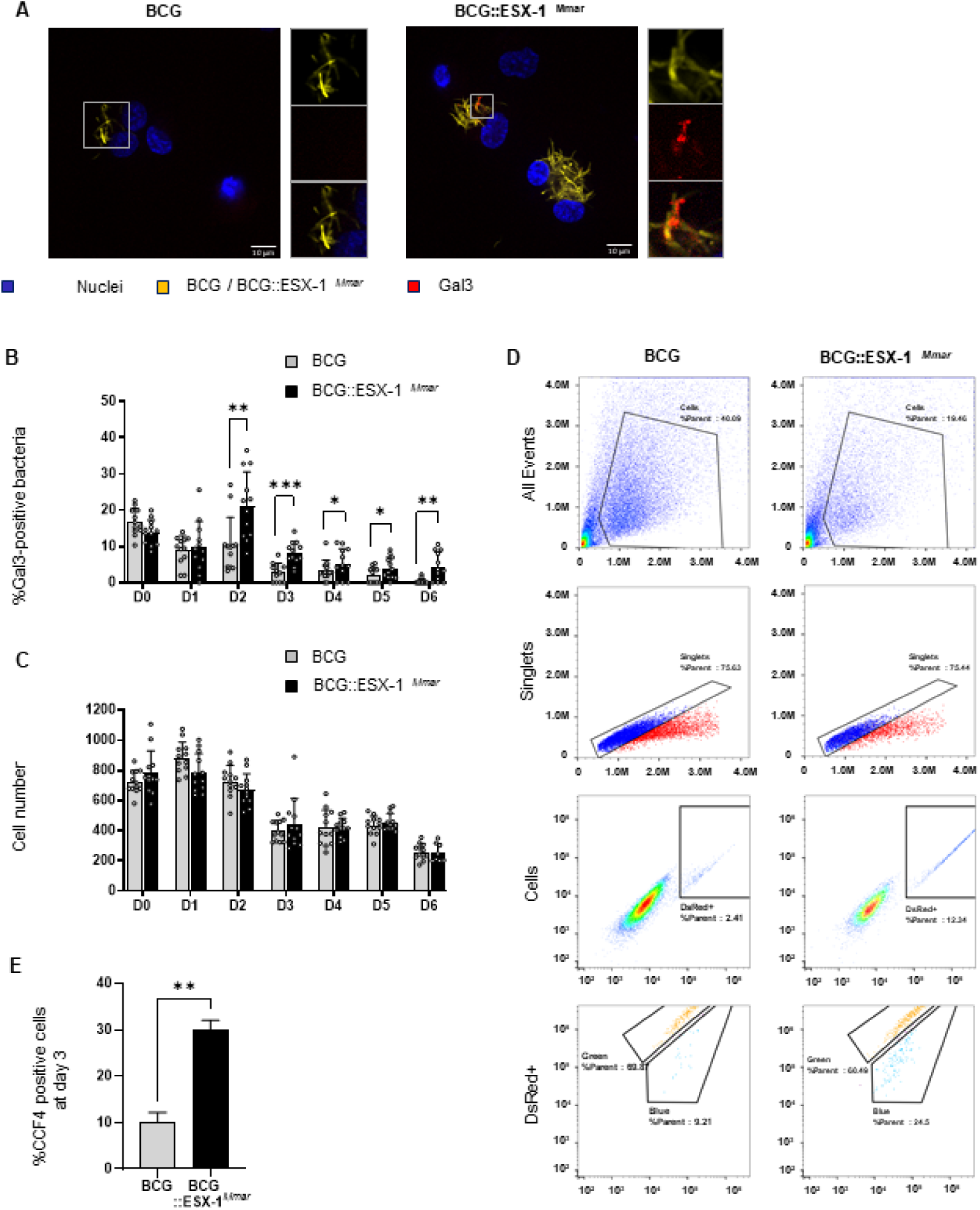
ESX-1–dependent phagosomal damage using BCG and BCG::ESX-1*^Mmar^*strains. THP-1 cells were grown in 384-well plates, infected with BCG pMRF1 and BCG::ESX-1*^Mmar^*pMRF1 (MOI = 2) for 6 days and imaged with confocal microscope. (A) Representative images of THP-1 (nuclei; blue) infected with BCG and BCG::ESX-1Mmar (yellow), showing bacterial colocalization of BCG::ESX-1*^Mmar^*with a Gal3 spot (red). Scale bar = 10 μm (60X). (B) Percentage of bacteria colocalizing with Gal3 spots over 6 days. BCG::ESX-1*^Mmar^* display a significantly higher rate of colocalization with Gal3 than BCG from day 2 post-infection (Mann-Whitney test, P≤0.05). (C) Quantification of total cell number under conditions of BCG and BCG::ESX-1*^Mmar^* infection. No significant difference is observed between the two bacterial strains (Unpaired t-test). (D) Flow cytometry gating strategy for BCG and BCG::ESX-1*^Mmar^*-infected samples. Singlet cells were first selected based on size (Forward Scatter, FSC) vs. granularity (Side Scatter, SSC) and on FSC-H vs. FSC-A, followed by separation of infected populations (DsRed+) displaying two distinct phenotypes based on blue and green fluorescence signals of the CCF4-AM staining. (E): Percentage of CCF4-positive (blue) infected cells at 3 days post-infection, which is significantly increased under BCG::ESX-1*^Mmar^* infection conditions than under BCG infection conditions (Unpaired t-test, P≤0.05).

To directly assess phagosomal rupture, we performed a CCF4-AM β-lactamase reporter assay by flow cytometry. This assay relies on a cytoplasmic FRET-based substrate that shifts fluorescence from green to blue upon cleavage by β-lactamase, indicative of bacterial access to the host cytosol following phagosomal membrane disruption [8]. Notably, cells infected with BCG::ESX-1*^Mmar^* exhibited a significantly higher proportion of blue-shifted signal compared with both BCG-infected and non-infected cells, consistent with increased phagosomal rupture in the presence of a functional ESX-1 system (Figures 2D-E).

Taken together, these results support a direct association between ESX-1–mediated secretion and phagosomal membrane disruption. However, despite the increased phagosomal rupture observed in BCG::ESX-1*^Mmar^*–infected cells, no significant differences in cell viability were detected between conditions over the six-day infection period. These findings indicate that ESX-1–dependent phagosomal rupture alone is insufficient to induce macrophage mortality in THP-1 cells, suggesting that additional virulence determinants are required to drive host cell death during mycobacterial infection.

### Galectin-3 loss results in an increased replication of H37Rv

We next investigated the functional contribution of Gal3 to the intracellular infection phenotype of *Mtb*. To this end, THP-1 macrophages were transfected with an siRNA targeting LGALS3, thereby suppressing Gal3 expression through mRNA silencing using protocols that were developed previously in our laboratory [26, 27]. Based on the previous observations, and to ensure data robustness in the context of infection-associated cell loss and the decrease in efficiency of the siRNA silencing, subsequent analyses were focused on the first three days post-infection. It allowed reproducible assessment of Gal3 distribution and its silencing, as well as maintaining sufficient cell numbers for quantitative analysis. Infection phenotypes were studied using live H37Rv or HK H37Rv, with a non-targeting scramble siRNA as a negative control, and siVPS18, as a positive control, as previously reported [15].

At this time point, LGALS3 silencing resulted in an approximately 60% reduction in Gal3 expression (Figures 3A-B). Consistent with this knockdown, Gal3 spot colocalization with H37Rv was significantly reduced in LGALS3-silenced cells, in contrast to the sharp increase of the Gal3-H37Rv colocalization obtained by VPS18 silencing (Figure 3C). Despite the decrease in Gal3 recruitment, LGALS3-silenced cells exhibited a modest but significant increase in intracellular bacterial area compared with scramble controls at five days post-infection, a phenotype similar to that observed following VPS18 knock-down (Figure 3D). No evidence of cellular toxicity was detected under any of the conditions tested (Figure 3E).

**Fig 3.**
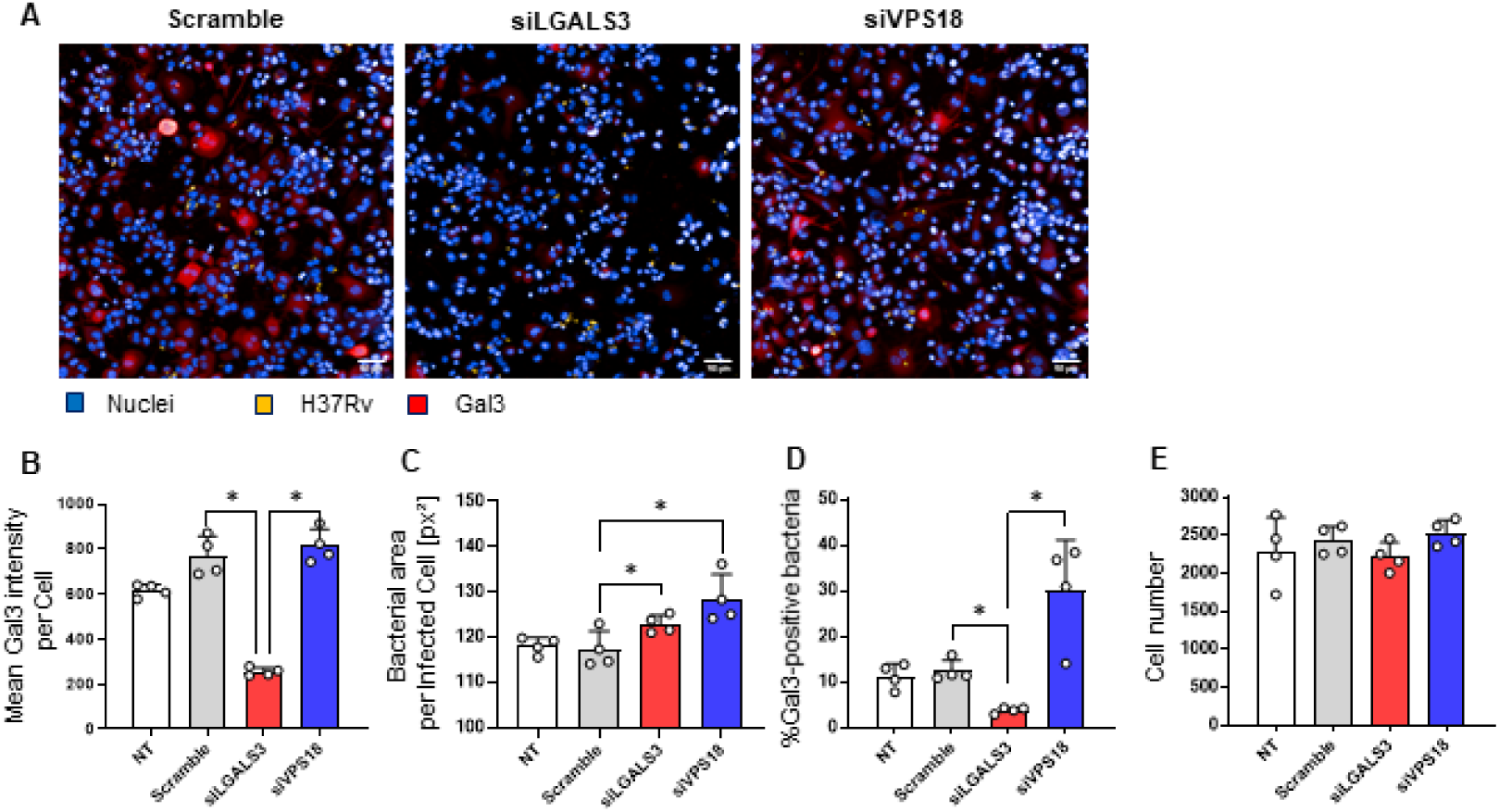
Functional impact of LGALS3 silencing on Galectin-3 levels and infection phenotypes. THP-1 cells were transfected with SiRNA in 384-well plates, infected with H37Rv pMRF1 (MOI = 1) for 3 days and imaged by automated confocal microscopy. (A) Representative images of THP-1 cells transfected with scramble control (left), siLGALS3 (middle) and siVPS18 (right) and infected. Scale bar = 50 μm (20X). (B, C, D, E) Quantitative image analysis of non-transfected (NT), scramble, siLGALS3 and siVPS18 conditions on day 3 post-infection. (B) Mean Gal3 fluorescence intensity. Gal3 signal is 4-fold reduced for siLGALS3-treated cells relative to scramble and siVPS18-treated ones (Brown–Forsythe ANOVA, P≤0.01). (C) Intracellular bacterial area (pixel²) per infected cells containing at least one bacterium that is positive for Gal3 (Gal3+, rupture) compared to cells infected with bacteria not colocalized with Gal3 spot (Gal3-, non-rupture). Bacterial area is significantly increased under both siLGALS3 and siVPS18 conditions by comparison with scramble condition (Unpaired t-test, P≤0.05). (D) Percentage of bacteria colocalizing with Gal3 spots. No significant difference is observed between NT and scramble controls. Bacteria and Gal3 colocalization is significantly decreased for siLGALS3-transfected cells and significantly increased for siVPS18-transfected ones relative to scramble condition (Mann-Whitney test, P≤0.05). (E) Total cell number for transfected and non transfected conditions. No significant difference is observed (Unpaired t-test).

Together, these findings suggest that Gal3 contributes to host defense mechanisms which restrict intracellular *Mtb* replication in THP-1 macrophages. Furthermore, these data confirm the role of VPS18 in limiting *Mtb* growth, likely through continuous regulation of phagosomal membrane integrity in the days following infection.

### No recruitment of Gal3 observed in human alveolar epithelial cells during the infection

To assess the cell-type specificity of the Gal3 associated phenotype observed in macrophages, we extended our analysis to primary human alveolar epithelial cells (HPAEpiC) over a 3-day infection period (Figures 4A-C). The proportion of HPAEpiC exhibiting Gal3 spots remained low and stable (approximately 4–5%) across all conditions, with no measurable variation in overall Gal3 signal intensity. No significant differences in cell numbers were detected between infected and non-infected conditions, in contrast to the pronounced cell loss observed in macrophage models (Figure 4B). Consistently, quantitative analysis showed no significant difference in bacterial burden between cells displaying Gal3 colocalization and those lacking detectable Gal3 recruitment, despite a modest increase in bacterial load over time (Figure 4D). No significant differences in cell numbers were detected between infected and non-infected conditions (Figure 4E).

**Fig 4.**
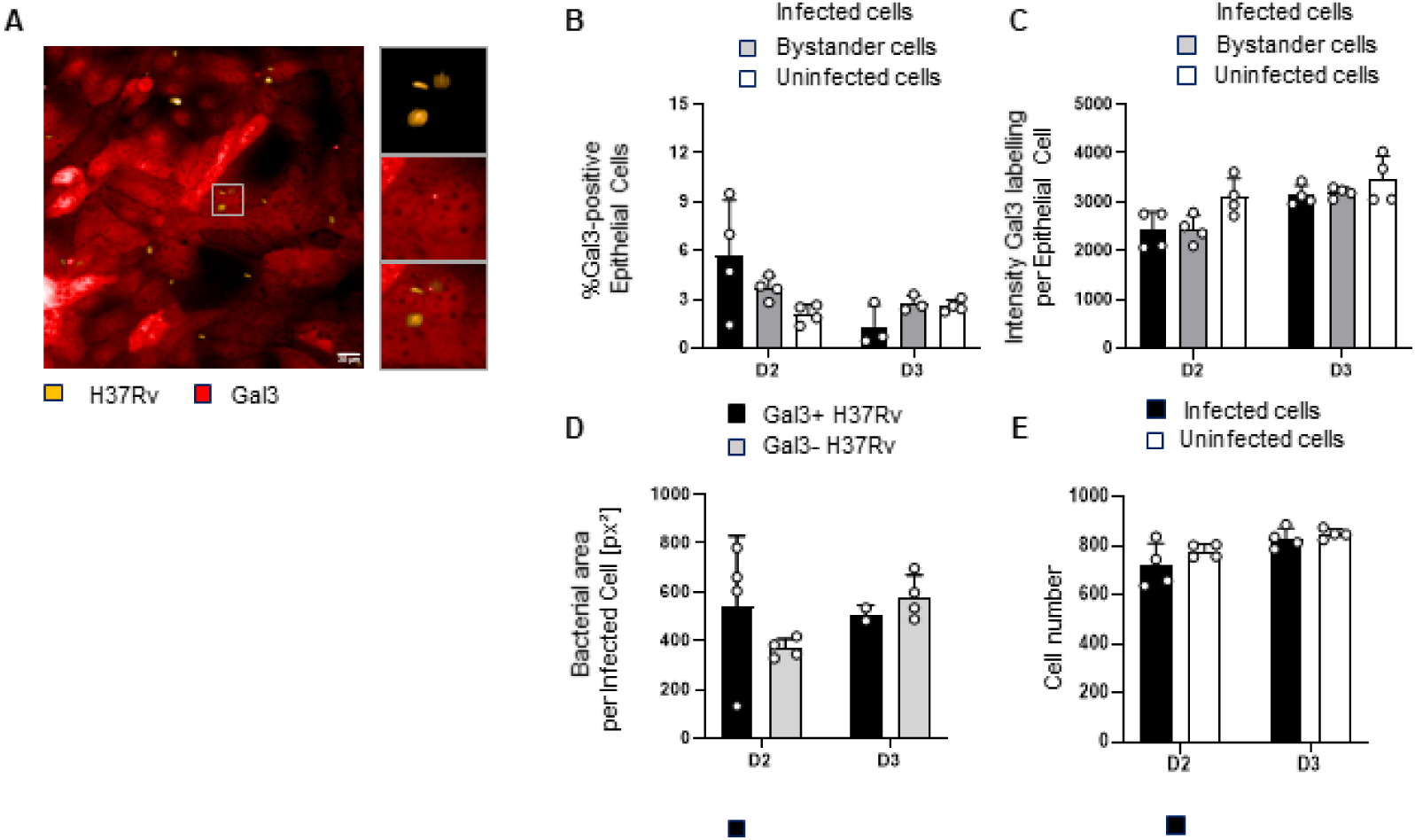
Infection phenotype and Galectin-3 distribution in human alveolar epithelial cells (HPAEpiC). HPAEpiC cells were grown in 384-well plates, infected with Mtb H37Rv pMRF1 (MOI = 5) for 3 days and imaged by automated confocal microscopy. (A) Representative image of HPAEpiC cells infected with live H37Rv pMRF1 bacteria (yellow) and labelled by immunofluorescence for Galectin-3 (Gal3; red). The focused image shows a bacterium which colocalized with a Gal3 spot. Scale bar = 20 μm (40X). (B, C, D, E) Quantitative image analysis of infected and bystander HPAEpiC cells infected with live H37Rv pMRF1 or uninfected over 3 days. (B) Percentage of infected, bystander and uninfected Gal3-positive HPAEpiC cells. No significant difference is observed between infected and bystander cells (Paired t-test) and between uninfected and bystander cells or uninfected and infected cells (Mann-Whitney test). (C) Mean Gal3 fluorescence intensity in infected, bystander and uninfected HPAEpiC cells. No significant difference is observed between cells infected with H37Rv pMRF1 and bystander cells (Paired t-tests) or uninfected conditions (Unpaired t-test). (D) Intracellular bacterial area (pixel²) per infected cells containing at least one bacterium that is positive for Gal3 (rupture) versus infected cells without colocalization of bacterium with Gal3 spot (non-rupture). The cells exhibiting colocalization with Gal3 or not show non significant difference of bacterial area from 2 days after infection. (Mann-Whitney test). (E) Total cell number for infected and non infected conditions. No significant difference observed between the conditions (Unpaired t-test).

Collectively, these data indicate that primary alveolar epithelial cells, in contrast to macrophages, do not exhibit substantial Gal3 redistribution or alteration in response to infection.

These findings highlight a marked cell-type specificity in the host response to *Mtb*, suggesting that Gal3-associated membrane damage and its downstream consequences are predominantly specific to macrophages.

### Similar phenotype of colocalization observed in an Alveolus-on-Chip model of infection

Initial time-course analyses of Gal3 dynamics in THP-1 macrophages prompted us to investigate its spatial distribution in a more physiologically relevant alveolar context. To do so, we used a human alveolus-on-chip (AoC) model that we recently set [22]. This organ on chip system recapitulates an alveolar barrier by co-culturing primary human alveolar epithelial cells (HPAEpiC) and integrating human macrophages (HuMacro) derived from peripheral blood mononuclear cells (PBMCs). This immunocompetent epithelium is interfaced with human lung endothelial cells (HuLECs) through a porous membrane, thereby mimicking the epithelial–endothelial 3D organization of the alveolar barrier (Figures 5A-B). This model was shown to support *Mtb* H37Rv infection over 1–5 days following apical inoculation at a multiplicity of infection (MOI) of 5, selectively targeting epithelial cells and macrophages.

**Fig 5.**
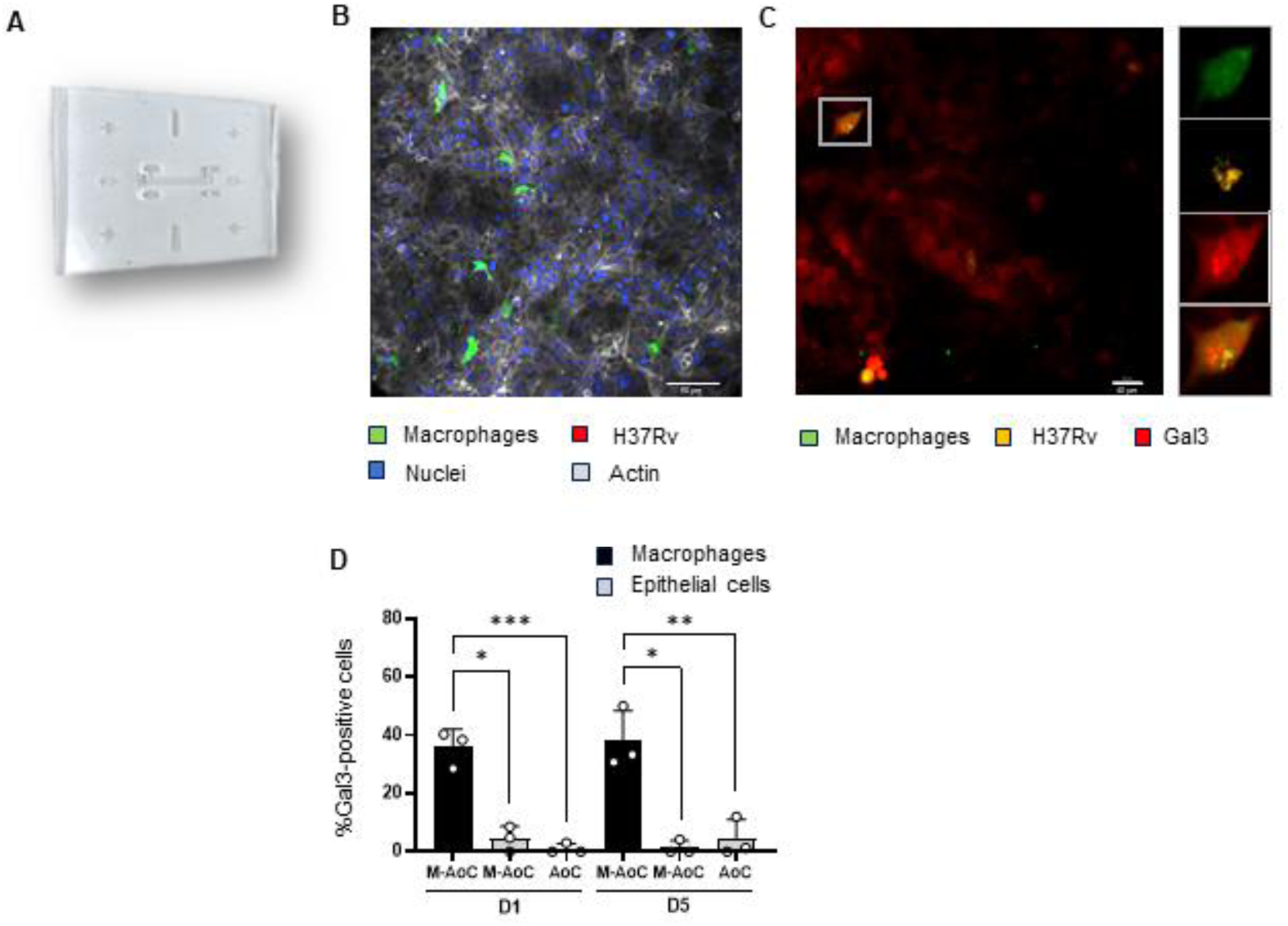
Human alveolus-on-chip (AoC) model and Galectin-3 responses to infection. HPAEpiC cells and human lung endothelial cells were seeded in in-house AoC devices, respectively in top and bottom channels, and grown over 7 days to reconstitute the alveolar interface. Human macrophages were introduced into the top channel on the day 7 (M-AoC) and then infected with Mtb H37Rv pMRF1 (MOI = 5) on day 8. M-AoC was imaged at 1 and 5 days post-infection by confocal microscopy and analyzed using a 3D image analysis software (IMARIS v10, Oxford Instruments). (A) Photograph of the in-house alveolus-on-chip (AoC) device. (B) Representative 3D-imaging of infected M-AoC at 1 day post-infection, used for the determination of H37Rv pMRF1 infection (red) inside the alveolar barrier model (nuclei, blue; actin, gray; macrophages, green). Scale bar = 50 μm (20X). (C) Representative immunofluorescence image of the alveolar barrier co-cultured with macrophages (CFSE labeled, green) and infected with H37Rv pMRF1 (yellow). The focused image highlights a bacterium within a macrophage colocalized with a strong Gal3 signal (red), consistent with phagosomal rupture. (D) Percentage of infected Gal3-positive HPAEpiC cells and macrophages in AoC and M-AoC models at days 1 and 5 post-infection. Infected macrophages exhibit significant higher rate than epithelial cells in M-AoC (Paired t-test) and epithelial cells in AoC models (Unpaired t-test, P≤0.05). Quantifications were obtained from pooled datasets comprising ∼10 infected macrophages and ∼40 infected epithelial cells per experiment.

Building on this framework, we examined Gal3 distribution in the AoC model under infection conditions (Figure 5C). Within the AoC environment, PBMC-derived HuMacro displayed a Gal3 distribution pattern closely resembling that observed in THP-1 macrophages cultured on plates. Gal3 was accumulated at sites of bacterial localization and approximately 30% of infected macrophages exhibited Gal3 spots colocalizing with H37Rv (Figure 5D). Similar to observations in 2D monolayer cultures, Gal3 spots were observed in only 4–5% of infected alveolar epithelial cells at either 1- or 5-days post-infection within the AoC environment, irrespective of the presence of HuMacro (Figure 5D).

Taken together, these results demonstrate the robustness and reproducibility of Gal3 immunostaining in the Alveolus-on-Chip model and confirm that Gal3 recruitment to *Mtb* occurs predominantly in macrophages rather than epithelial cells.

## Discussion

In this study, we investigated the long-term dynamics of phagosomal rupture and Gal3 during *Mtb* infection by combining a longitudinal imaging with genetic perturbation. While phagosome rupture is mainly studied during the early events of *Mtb* infection, our monitoring of *Mtb* infection over five days, using conventional in vitro and biomimetic alveolus on chip models, revealed an unexpected, sustained recruitment of Gal3 along *Mtb* intracellular infection, supporting the existence of distinct early and late phase of membrane damage. Importantly, these two phases differed in their dependence on bacterial viability and ESX-1 activity, suggesting a more complex mechanism than previously known. Furthermore, we observed that Gal3 depletion increased intracellular replication predominantly in macrophages, supporting a model in which Gal3 is not merely a passive marker of membrane damages but contributes differently in the host defense mechanism for restricting bacterial replication. Finally, the comparison of this process between macrophages and alveolar epithelial cells reveals a distinct phenotype and further supports a model where *Mtb* intracellular outcome is strongly influenced by its distinctive cellular niches within the alveolar environment.

While Gal3 was considered as a hallmark of early infection, usually few hours following bacteria endocytosis, our longitudinal imaging revealed that Gal3 colocalization with intracellular bacteria persists for several days, suggesting that phagosomal membrane damage is a sustained process and not restricted to early events. Interestingly, we observed that the level of Gal 3 recruitment was similar within the first few hours following infection with live or heat-killed bacteria. This suggests that the initial Gal3 recruitment is likely not solely the result of bacterial virulence activity but may partly reflect a host-driven process associated to phagocytosis membrane remodeling or a transient upregulation of Gal3, which had previously been linked to autophagy pathways [20]. Consistent with this interpretation, Gal3 spots were predominantly detected in infected cells rather than bystanders at early stages. In contrast, starting one day after infection, infected cells displayed significantly reduced overall Gal3 expression relative to neighboring cells, supporting the hypothesis that bacterial effectors actively modulate Gal3 levels during intracellular residence. This role is consistent with previous studies describing Gal3 as a host-protective factor in other infections with *Toxoplasma gondii* and *Helicobacter pylori* [28, 29]. Interestingly, the sustained Gal3 recruitment within the “late” phase was associated with active bacteria and ESX-1 competent strains, further supporting the central role of ESX-1 for mediating membrane damage during *Mtb* infection. These results are consistent with previous models where ESX-1 and PDIM are essential for cytosolic bacterial access during *Mtb* infection.

Beyond its use as a marker of membrane damaged associated to phagosomal rupture, several evidences provide support that Gal3 actively contribute in host cell response. While the precise mechanism remains to be determined, the depletion of Gal3 led to an increase of intracellular bacterial replication, hence strongly supporting a protective role of Gal3 in *Mtb* infection. Interestingly, depletion of VPS18 produced an opposite phenotype for Gal3 recruitment while increasing bacterial growth, reinforcing the importance of membrane integrity as a protector against infection [15]. This opens new avenues for better understanding the relationship between membrane integrity, autophagy and inflammatory signaling in this process.

An important result in our study is the observation that the recruitment of Gal3 is predominant to macrophages. While alveolar epithelial cells could be infected, we observed a different phenotype, including a minimal recruitment of Gal3 and no modulation of Gal3 expression during infection. This difference in phenotype suggests that *Mtb*-host interaction may fundamentally differs between cell type infected and highlights the supportive conditions for sustaining specific distinctive cellular niche. Importantly, these phenotypes were reproduced in conventional in vitro cell culture and in biomimetic conditions using a human alveolus on chip [22]. The consistency observed between our models validate the robustness of this Gal3 associated phenotype. While the exact mechanism remains to be determined, our results are consistent with previous models proposing epithelial cells as a reservoir of infection rather than active places of replication and dispersion by *Mtb* [4].

Overall, our results identify Gal3-associated membrane damage as hallmark of *Mtb* infection in macrophages and reveal that phagosomal damage dynamics and intracellular bacterial replication are actively organized temporally. In contrast to previous studies in *Mtb* and other pathogens[12, 14, 30], our study reveals that *Mtb* phagosomal membrane damage is a sustained, rather than transient, process. More broadly, our framework integrating longitudinal imaging, genetic perturbation and human biomimetic lung on chip model offers new opportunities for exploring the mechanism governing phagosomal membrane damage toward the identification of new host-directed therapeutic strategy targets during *Mtb* infection.

## Experimental Procedures

### Cell culture and differentiation

THP-1 cells (ATCC; Cat#TIB-202), a human monocytic leukemia cell line, were maintained in suspension in complete RPMI-1640 medium supplemented with L-glutamine and 10% heat-inactivated fetal bovine serum (FBS; 56°C, 30 min). THP-1 cultures were maintained at 37 °C with 5% CO₂ and resuspended every 3–4 days in fresh medium at 2 × 10⁵ cells/mL, as previously described [27]. 50 µL of cells at 6 × 10⁵ cells/mL, supplemented with 50 ng/mL phorbol 12-myristate 13-acetate (PMA), were plated in a 384-well plate (Greiner µCLEAR Black; Cat#781091) and incubated for 72 h before infection, inducing macrophage-like differentiation and adherence.

Endothelial cells (HuLEC-5a; Cat#CRL-3244; ATCC), primary human alveolar epithelial cells (HPAEpiC; Cat#H-6053; Cell Biologics), and human macrophages derived from peripheral blood mononuclear cells (HuMacros) were cultured, as previously described [22]. Briefly, HuLEC-5a cells were grown in MCDB 131 medium (Gibco) supplemented with 10% heat-inactivated FBS, 10 mM L-Glutamine, 1 µg/ml Hydrocortisone (Cat# H0888; Sigma-Aldrich), 10 ng/ml recombinant human Epidermal Growth Factor (Cat# AF-100-15; PeproTech) and 1% Penicillin/Streptomycin in 5% CO_2_ at 37°C. HPAEpiC were grown in a treated culture flask pre-coated with gelatin-based coating solution (Cat# 6950; Cell Biologics) and incubated in complete Epithelial Cell growth medium comprising supplements (Cat# H-6621; Cell Biologics) at 37°C, 5% CO_2_. Cryo-preserved vials, containing at least 0.5 × 10^6^ cells at P4 per ml were thawed and used for seeding. 50 µL of cells at 1 × 10^6^ cells/mL were plated in a 384-well plate and incubated for 24 h before infection. Human monocytes were previously isolated from peripheral blood mononuclear cells (PBMCs) from buffy coats from anonymized donors (provided by Regional French Blood Company, France), following described protocol (45). Cryo-preserved monocytes were thawed and differentiated into human macrophages (HuMacro) by culture for 7 days in RPMI medium (Gibco) supplemented with 10% FBS, 1% Penicillin/Streptomycin and 40 ng/mL recombinant human macrophage-colony stimulating factor protein (hM-CSF, Cat# 130-096-491; Miltenyi).

### siRNA transfection of THP-1 cells in 384-well plates

siRNA transfection was concomitantly performed with the treatment of THP-1 with PMA into 384-well plates over the 72-h period before infection. The assays were done using SMARTPool ON-TARGETplus siRNA from Horizon Discovery of LGALS3 (Cat#L-010606-00-0005), VPS18 (Cat#L-013178-00-0005), and Non-targetting (Scramble) as a negative control (Cat#D-001810-10). siRNAs were first pre-dispensed into wells prior to cell seeding, as previously described (45). Briefly, 250 nL of siRNA pre-diluted in 1× siRNA Buffer (Cat#B-002000-UB-100; Horizon Discovery) per well were deposited using an automatic liquid handler (Echo 550, Labcyte). Subsequently, 10 µL of Lipofectamine RNAiMAX (Cat#13778030 ; Thermo Fisher Scientific), diluted 1:100 in DPBS without Ca²⁺ and Mg²⁺ (DPBS -/-), per well, were added to mediate transfection. The siRNA–lipid complexes were allowed to form for 30 min at room temperature before cell addition. THP-1 cells were then adjusted to 7.5 × 10⁵ cells/mL and 40 µL of suspension was dispensed per well, ensuring uniform seeding density across conditions.

### Bacterial strains and culture conditions

Recombinant strains of DsRed-expressing BCG Pasteur reference strain (BCG), BCG Pasteur strain expressing the ESX-1 secretion system from *Mycobacterium marinum* (BCG::ESX-1*^Mmar^*), and virulent *Mtb* H37Rv were transformed by pMRF1, a derivative of pMV261 carrying a kanamycin resistance cassette, allowing the expression of the red fluorescent protein DsRed [31]. These strains were cultured in Middlebrook 7H9 medium (Difco) supplemented with 10% oleic acid-albumin-dextrosecatalase (Difco), 0.2% glycerol (Euromedex), 0.05% Tween 80 (Sigma-Aldrich), and 25 µg/ml kanamycin (Sigma-Aldrich) and 25 µg/mL hygromycin B (Sigma-Aldrich), for DsRed-expressing strains and ESX-1*^Mmar^* -expressing strain respectively. Culture were grown for up to 14 days before being used for the infection assay, to reach the exponential phase of bacterial growth, as previously described [27]. Bacterial growth was monitored spectrophotometrically by optical density (OD) measurement.

### *Mtb* infection in 384-well plate

*Mtb* culture grown to exponential phase (OD600 of 0.5) was used as the standardized condition for infection assays and prepared as previously described [27]. Briefly, after supernatant removal by centrifugation, the cell pellet was washed two times with DPBS -/- and resuspended in complete media. Clumped mycobacteria were removed by centrifugation at 800 rpm for 2 min. And a single cell suspension was obtained via filtration through a 10 µm cell pluristrainer (Cat# 43-50010-01; pluriSelect). Optical density at 600 nm was measured.

The single-bacteria suspension of H37Rv-pMR1 was diluted to obtain a multiplicity of infection (MOI) of 1 and 5 for infection of THP-1 and HPAEpiC, respectively. And the suspensions of BCG and BCG::ESX-1*^Mmar^* were adjusted to MOI 2 for infection of THP-1. Heat-killed controls were prepared from the H37Rv pMRF1 suspensions by heat treatment at 90 °C for 30 min. Prior to infection, 50 µL of medium was removed from each well and replaced with 50 µL bacterial suspension prepared at the appropriate MOI, then incubated at 37 °C, 5% CO₂ for 2 h (THP-1) or 4 h (HPAEpiC). Wells were subsequently washed three times with 50 µL sterile medium. For epithelial cells, amikacin was added to 50 µg/mL for 1 h to eliminate extracellular bacteria, followed by two additional washes. Cells were then incubated for up to 5 days before downstream assays. Bacterial inocula were validated by colony-forming unit (CFU) assays. Serial 10-fold dilutions were spotted (2 µL) onto 7H11 agar plates with hygromycin (ESX-1 strains) or without antibiotics (BCG and H37Rv). After drying, plates were incubated at 37 °C with 5% CO₂ for ≥20 days before manual colony counting. Absence of colonies in heat-killed controls confirmed successful inactivation.

### Preparation of human alveolus-on-chip and *Mtb* infection

To generate the human alveolus-on-chip (AoC), HuLEC-5a and HPAEpiC cells were seeded into the 3DP-µLung microdevices, composed by two distinct fluidically independent top and bottom channels separated horizontally by a polyester porous membrane, as previously described [22]. Briefly, both channels of 3DP-µLung were coated overnight at 37°C with 200 μg/ml Collagen IV (Cat# C5533; Sigma-Aldrich) and 30 μg/ml Fibronectin from human plasma (Cat# 356008; Corning). 10 µL containing 1 × 10^5^ of pre-cultured HuLEC-5a cells were seeded in the bottom channel on next day. Chips were flipped and placed on chip cradles for 4 h at 37°C, 5% CO_2_. After to be flipped back and washed-out unattached cells, chips were maintained overnight under static conditions. On day 1, HPAEpiC were thawed and 20 µl of 5 × 10^4^ cells were seeded in the top channel and incubated for 4 h at 37°C. Before wash, chips were incubated statically overnight. The seeded 3DP-µLung were connected to microfluidic circuit actuated by a peristaltic pump (Ismatec, Switzerland), regulating media flow (60 µl/h) in both channels. Apical and basal channels were perfused with cell-specific growth media, and epithelial medium was supplemented with 1 μM dexamethasone (Cat# D4902; Sigma-Aldrich) to promote surfactant production and formation of tight junctions. Chips were cultured under flow conditions for 4 days at 37°C, 5% CO_2_. On day 6, the apical channel medium is removed, to establish an air-liquid interface (ALI). Chips were fed through the bottom channel for one day. AoC with ALI were detached from microfluidic system on day 7 and kept with a liquid-liquid interface (LLI) for AoC experiments. For AoC co-cultured with human macrophages (M-AoC experiments), HuMacro were labeled on day 7 with CellTrace CFSE solution at a final concentration of 10 µM (Cat# C34570; Invitrogen), following the steps described in the manufacturer’s instructions. 20 µl containing 5 × 10^3^ CFSE-labeled macrophages (corresponding to 10% epithelial cells) were injected into the top channel of the chips which were kept in static condition overnight at 37°C, 5% CO_2_. 8-days old AoC or M-AoC prepared as described above were washed before infection with 20 µl of DsRed-expressing *Mtb* at an estimated MOI of 5 into the epithelial channel. Chips were incubated in static condition for 4 h at 37°C, 5% CO_2_ to allow *Mtb* infection on the epithelial face; Channels were washed with epithelial and endothelial medium and treated with amikacin for 1 h at 37°C to kill extracellular bacteria. The AoC was washed two fold more and returned to LLI in static condition for 1 up to 5 days more at 37°C, 5% CO_2_, respectively, corresponding to day 9 and day 13. Serial 10-fold dilutions of the inoculum were plated on 7H11 plates and CFU counted after 2 weeks of incubation at 37°C, to determine the infectious dose.

### Galectin-3 immunostaining

After 1 to 5 days of infection, cells were fixed and immunolabelled for Gal3 detection. For 384-well plates, the culture medium (50 µL per well) was replaced with 50 µL of 4% paraformaldehyde solution (PFA) containing Hoechst at 10 µg/mL. Plates were incubated at room temperature (RT) for 30 min, followed by a single wash with DPBS -/-. Wells were incubated with 50 µL of 50 mM NH₄Cl for 2 mins at RT to quench autofluorescence, then washed three times with DPBS -/-.

For the AoCs, 4% freshly diluted PFA in DPBS with Ca^2+^ and Mg^2+^ (DPBS +/+; Gibco) was injected in two channels by inlets and the AoC devices completely immersed in PFA ON at RT. Following two washes with DPBS +/+, 50 mM NH_4_Cl solution was used for 30 min at RT and rinsed twice. Fixed AoC could be stored at 4°C.

Cells were then permeabilized using 0.3% Triton X-100 in DPBS for 5 and 15 min, in wells and inside AoCs respectively, and washed three additional times with DPBS -/-. Non-specific binding sites were blocked by incubation with 1% BSA in DPBS -/- (PBS-BSA 1%) for 30 min in wells under stirring, and for 1 h in AoC with 20 µg/mL Hoechst and Phalloidin A647 (1/400 dilution; A22287, Thermofisher), at RT, followed by 2 washings.

Plates and AoCs were incubated with the AlexaFluor 647-coupled anti-galectin-3 antibody (AF647-Gal3; BioLegend; Cat#125408), diluted 1:250 and 1:200, respectively, in 1% PBS-BSA, and moreover with the AlexaFluor 488-coupled anti-CD68 antibody (AF488-CD68; Santa Cruz; Cat# sc-20060 AF488) diluted 1:50 for AoCs, overnight at 4°C. The following day, wells and channels were washed twice, and kept at 4°C until imaging.

### CCF4-AM staining

CCF4-AM (Invitrogen, Cat#K1095) labelling experiments were carried out on 24-well plates, using an equivalent cell concentration of THP-1 cells by seeding 800 µL of cells. Four days after infection with BCG and BCG::ESX-1*^Mmar^*, CCF4-AM labelling was performed on live cells. The labelling mixture was prepared in EM buffer (120 mM NaCl, 7 mM KCl, 1.8 mM CaCl₂, 0.8 mM MgCl₂, 5 mM glucose, 25 mM HEPES, pH 7.3) and contained 8 µM CCF4-AM and 2.5 µM probenecid (Sigma-Aldrich ; Cat#P8761-25G). For each well, the mix consists of 20 µL of solution B, 3.2 µL of CCF4-AM (1 mM, prepared in DMSO and stored at -80°C) and 376.8 µL of EM buffer supplemented with probenecid.

Before adding the mix, the cells are rinsed twice with 800 µL of EM buffer. The CCF4 mixture is then added (400 µL/well), followed by incubation at room temperature in the dark for 1 hour for THP-1 cells. After incubation, the cells are rinsed once with EM buffer.

### Flow Cytometry

After the CCF4 staining, the buffer is removed, followed by the addition of 100 µL of 0.25% trypsin-EDTA (Gibco ; Cat#25200-056), and left for 5 minutes to allow the cells from each condition to detach. Each condition is then transferred to Eppendorf tubes before adding 300 µL of EM buffer. Approximately 30000 events per sample were then acquired on an Aurora CS flow cytometer (Cytek Bioscience), at a flowrate of 2000 events/second with a 85µm nozzle. Data were analyzed using SpectroFloCS (v1.2.2, Cytek) and FlowJo 10 (v10.10.0, Tree Star) software. Successive gatings were made to isolate singlet population of infected cells (DsRed positive) from non-infected cells (DsRed negative), and then separate cells displaying CCF4 green fluorescence (V3 detector, 451-466 nm) or CCF4 blue fluorescence (V7 detector, 533-590 nm). By the principle of the CCF4 FRET assay, cells expressing a ratio of blue/green fluorescence >1 were considered positive to phagosomal rupture.

### Image acquisition and analysis

Images were acquired using an automated confocal fluorescence microscope (InCell Analyzer 6000, GE Healthcare) equipped with a 20x objective for experiments performed in 384-well plates. Excitation was provided by lasers at 405, 561, and 640 nm, enabling simultaneous detection of nuclei, bacteria, and Gal3 immunostaining. For each well, six independent fields were acquired. Images were subsequently analyzed using Columbus image analysis software (v2.9.1, PerkinElmer), and values from the six fields were averaged to generate a single measurement per well.

Images from the automated confocal microscope were analyzed using multi-parameter scripts developed using Columbus system (version 2.3.1; PerkinElmer). Segmentation algorithms were applied to input images to detect nuclei, bacteria and Gal3 spots. Briefly, the host cell segmentation was performed using Hoechst and Gal3 far red signal intensities, corresponding to the nucleus and cytoplasm, respectively. The signal of DsRed *Mtb* H37Rv pMRF1 were used for the bacterial segmentation. Subsequently, the population of infected cells was determined, as the ratio of cells containing at least one detectable bacterium to the total number of cells per well, and the increase of intracellular *Mtb* area, corresponding to intracellular mycobacterial replication, was calculated, as intracellular *Mtb* area with number of pixels. Gal3-positive spots were defined as distinct regions with high fluorescence intensity that exceeded the average cytoplasmic Gal3 signal by at least twofold, a threshold indicative of membrane damage. For each intracellular bacterium, a region of interest with a 4-µm radius was defined, and Gal3 spots within this area were classified as phagosomal rupture events. The proportion of bacteria associated with Gal3 recruitment was then determined for each condition, called “Gal3-positive bacteria”. Moreover, Gal3 spots distributed across the entire cellular surface were identified in both infected and uninfected cells, allowing discrimination between bacteria-associated Gal3 recruitment and physiological or non-specific cellular processes.

AoCs were imaged using a spinning disk confocal microscope (CSU-W1 LiveSR, Gataca Systems coupled to Nikon) equipped with a 20× dry Nikon objective.

AoC samples were stained with Hoechst 33342 to label nuclei, AF488-conjugated CD68 to identify macrophages, and AF647-conjugated Gal3. H37Rv-pMRF1 fluorescence was also detected.

Higher-resolution images were obtained using a laser scanning confocal microscope (LSM 880, Zeiss) with a 60× oil immersion objective (NA 1.4).. The acquisitions were made on 3 independent regions of the AoC (corresponding to inlet, center, and outlet regions). Z-series of optical sections were taken from the endothelial cell layer on the lower face of the porous membrane into the bottom channel up to the top of the epithelial cell layer into the top channel. The images were then analyzed using Imaris 3D microscopy image analysis software (v11.0, Oxford Instruments), and the datas of each fields were then averaged to generate a single measurement per AoC. Nuclei (Hoechst-labeling) and H37Rv-pMRF1 bacteria (DsRed) were detected, using a volume and a spot detection, respectively, based on fluorescence intensities on the 405 nm and 561 nm channels. The cytoplasm was detected using the AF647-Gal3-labeling. And the AF488-CD68 staining was used to identify the macrophage population by volume detection. For each bacteria, the ratio between the intensity of the Gal3 staining on the bacteria and the mean intensity of Gal3 in the cell cytoplasm was calculated. Because of the low image resolution, the ratio of 1.5 was selected to determine the Gal3-positive bacteria. The percentages of infected cells positive for Gal3 were then calculated for the macrophage and the epithelial cell populations.

## Statistical Analysis

Graphing and statistical comparison of the data were performed using Prism 10.4.1 (GraphPad Prism Software). Two-group comparison was assessed using an unpaired t-test for independent studied conditions; and a paired t-test was used to compare infected and bystander cells in the same well. When the requirement for the previous tests weren’t met, non-parametric tests were done instead, such as Mann-Whitney for unpaired t-test, and Wilcoxon for paired t-test. One-Way Anova was used to compare the cell numbers. Each presented experiment was repeated at least 3 times, with 3 biological replicates for the AoC and flow cytometry up to 6 biological replicates for experiments on plate. Data are presented as mean values ± standard deviation (SD) unless otherwise noted in the figure’s legends. Values of p ≤ 0.05 were considered to be statistically significant.

**Table 1.** List of siRNAs and their accessions hit used for the transfection assay

| SiRNA (ON-TARGETplus) | Cat# |
| --- | --- |
| Non-targeting Pool (Scramble) | Cat#D-001810-10-20 |
| Human LGALS3 (NM_001357678, NM_002306, NR_003225) | Cat#L-010606-00-0005 |
| Human VPS18 (NM_020857, XM_011521843) | Cat#L-013178-00-0005 |

**Table 2.**
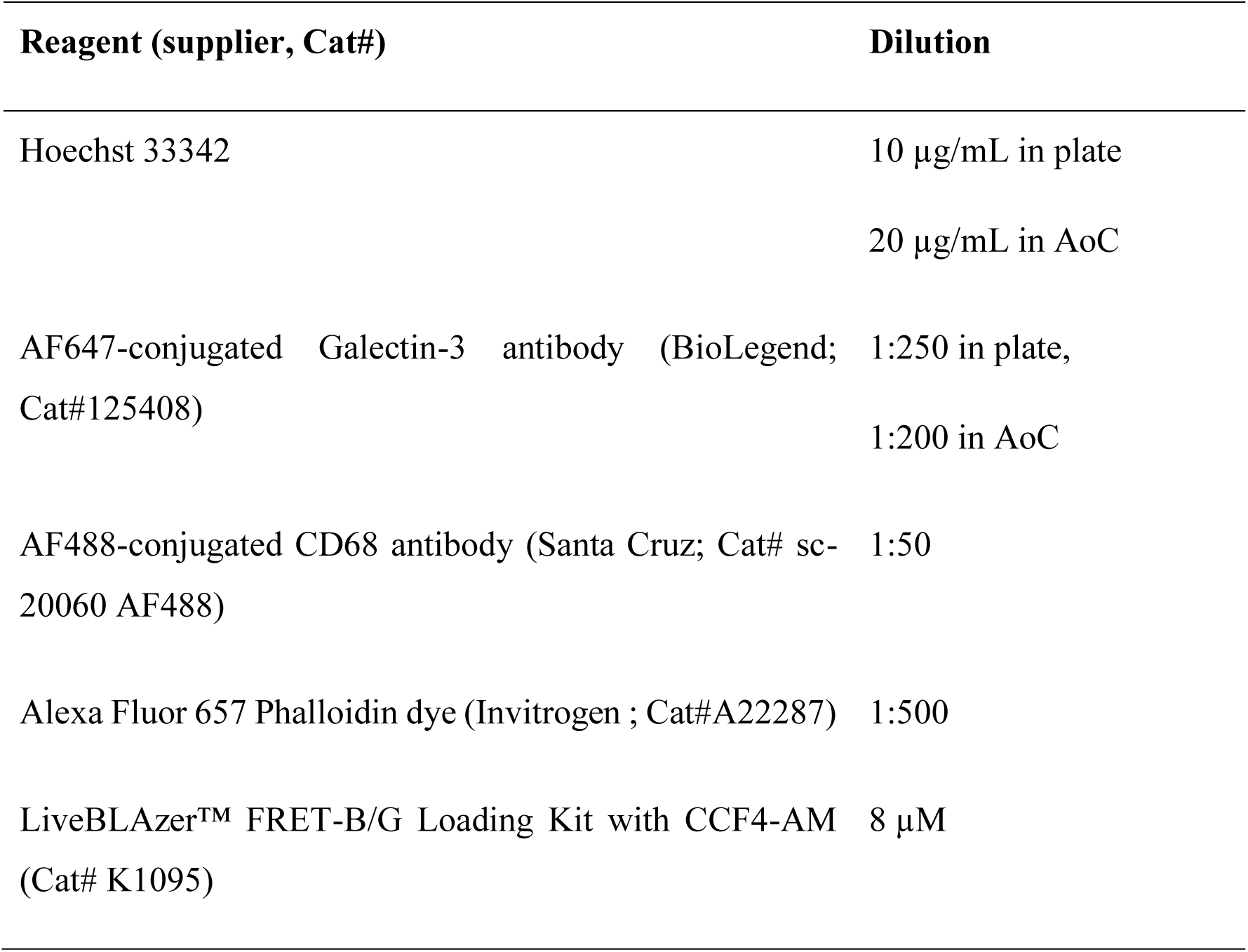
List of dyes and antibodies used for immunofluorescence staining

| Reagent (supplier, Cat#) | Dilution |
| --- | --- |
| Hoechst 33342 | 10 µg/mL in plate<br>20 µg/mL in AoC |
| AF647-conjugated Galectin-3 antibody (BioLegend; Cat#125408) | 1:250 in plate,<br>1:200 in AoC |
| AF488-conjugated CD68 antibody (Santa Cruz; Cat# sc-20060 AF488) | 1:50 |
| Alexa Fluor 657 Phalloidin dye (Invitrogen ; Cat#A22287) | 1:500 |
| LiveBLAzer™ FRET-B/G Loading Kit with CCF4-AM (Cat# K1095) | 8 µM |

## Acknowledgements

The authors thank Marc Settelen, Alexandre Vandeputte, Ines Leleu, Julien Lombard, Celine Wichlacz, Ruben Hartkoorn, and Jonathan Chatagnon for insightful discussions. They are also grateful to Cyril Gaudin, Eik Hoffmann and Manoly Wacheux for assistance with laboratory access. They thank Amine Pochet, Joan Fine, and Axelle Grande for help with some of the experiments. We thank the ARIADNE-Criblage platform (UMS2014-US41 PLBS) for providing access to the high-content automated microscope. The study benefited from valuable advice provided by Jost Enninga, Oana Dumitrescu, and Frank Lafont.

This work was supported by INSERM, the Agence Nationale de la Recherche (ANR-18-JAM2-0002, 20-AMRA-0005, and ANR-22-CE18-0021-01), the Agence Nationale de Recherche sur le SIDA et les hépatites virales - Maladies Infectieuses Emergentes (ANRS-MIE No. ANRS0521 and ANRS-0655), and the Institut Pasteur (PTR 430 and PTR 22-16). This project was co-founded by the European Union through the European Regional Development Fund (FEDER/ERDF) within the framework of the Contrat de Plan Etat-Ŕegion (CPER) 2021-2027 for the Hauts-de-France region. Y.D. was the recipient of a fellowship from the Agence Régionale de Santé (ARS). This project has received funding from the Innovative Medicines Initiative 2 Joint Undertaking (JU) under grant agreement No. 853989 (ERA4TB). This research is supported by Atip-Avenir funding, MEL “Accueil de talents” funding. French government under the France-2030 program, the University of Lille and the Lille European Metropolis (MEL) are thanked for their funding and support for the project R-CDP-24-007-MOSAIC granted to AG.

The funders had no role in study design, data collection and analysis, decision to publish, or preparation of the manuscript. The authors declare that they have no competing interests.

## Author contributions

Y.D designed, and performed the 2D experiments and data analyses. N.D. managed AoC experiments and performed maturation steps to generate AoC in BSL2 environment. Y.D. cultured the H37Rv strain and performed infection experiments in BSL3 environment. Y.D. analyzed the images using IMARIS software. E.B. performed the experiments using BCG and BCG::ESX-1*^Mmar^*strains. A.B. fabricated the microfluidic chips. M.S. helped develop the immunostaining assay. A.M. and R.S. helped for the correction of the article and gave valuable insights for the experiments. O.M. helped develop the flow cytometry assay. S.D. and E.W. participated in image analysis.

P.B. and A.G performed the conceptualization, funding acquisition, and supervision.

Y.D.; N. D.; P.B. and A.G. wrote the manuscript. All authors read, reviewed, and approved the final manuscript.

---

**Supp Fig 1.**
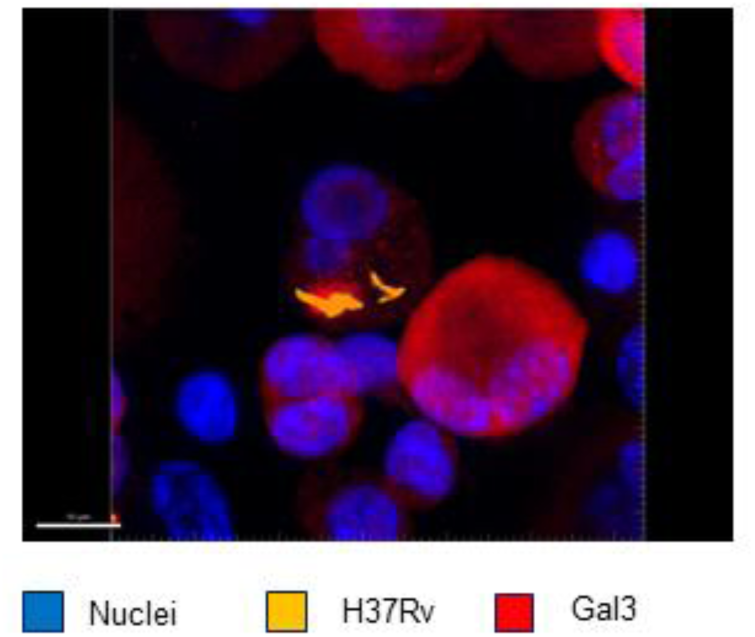
3D animated picture of an infected cell stained by anti- Galectin-3 antibody. 3D imaging of Gal3 spots colocalized with Mtb H37Rv pMRF1 (MOI 1; yellow) inside THP-1 cells (nucleus; blue) at day 3 post-infection, using 3D image analysis software (IMARIS v10, Oxford Instruments). Scale bar = 4-10 μm (60X).

**Supp Fig 2.**
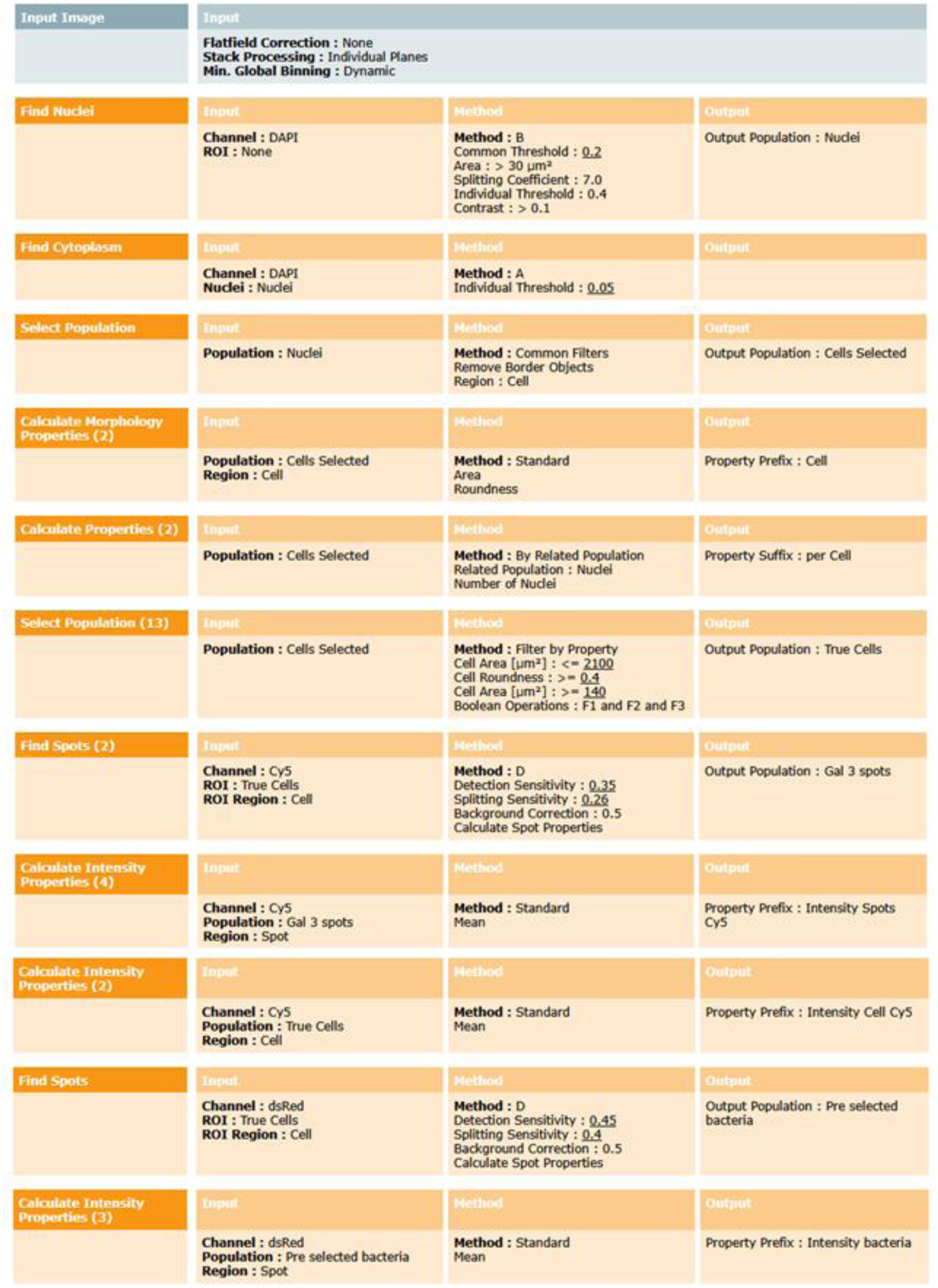
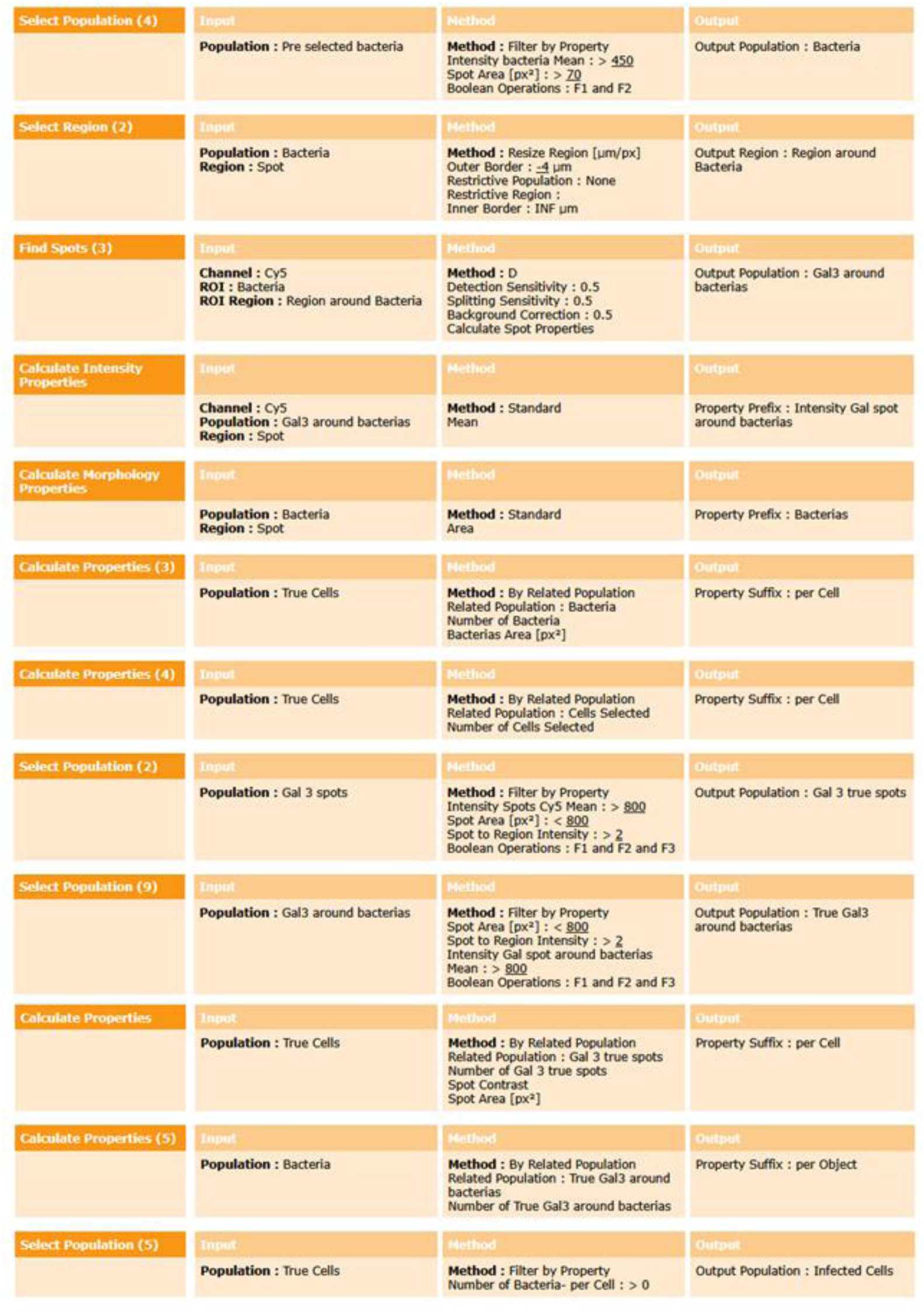
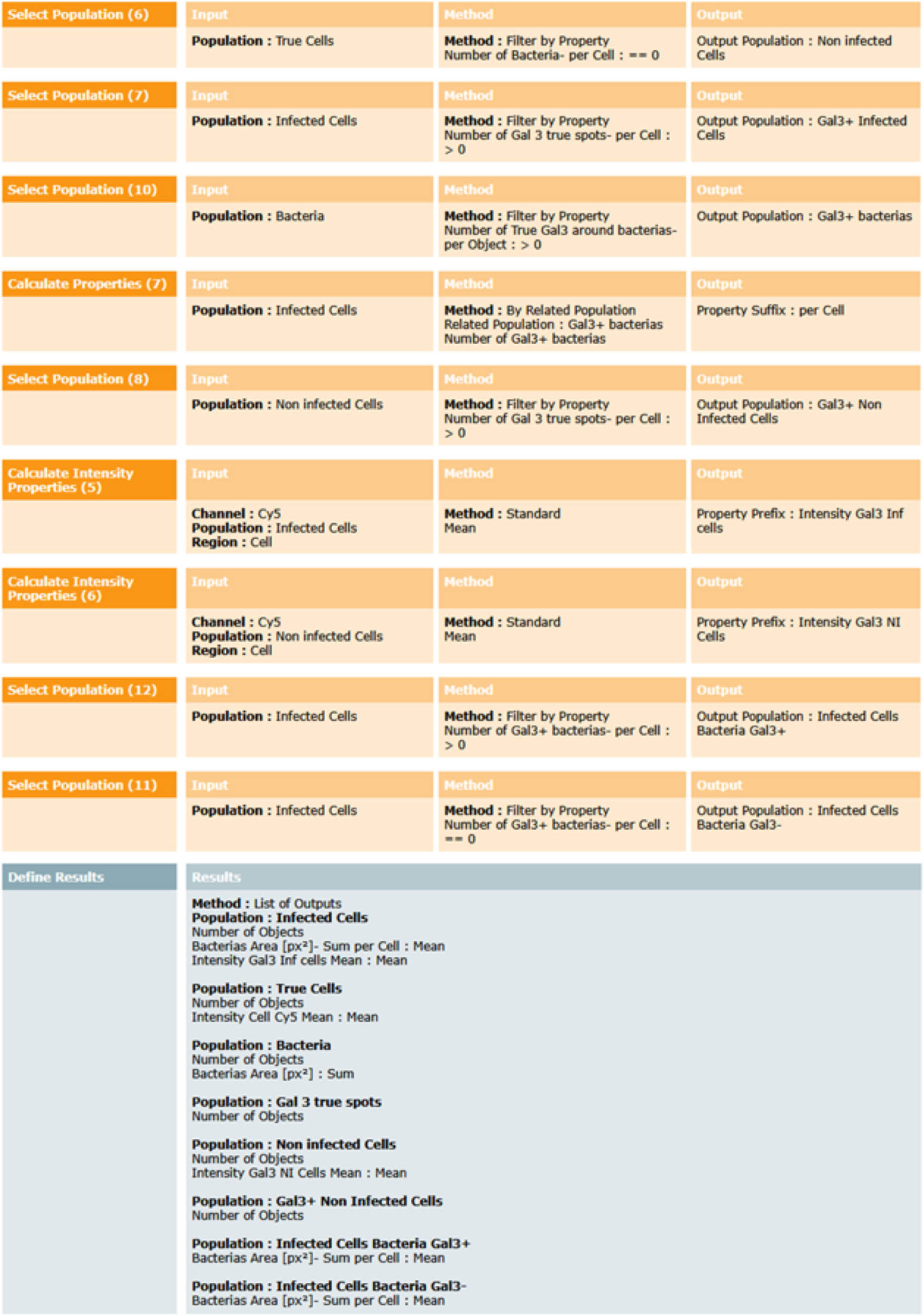
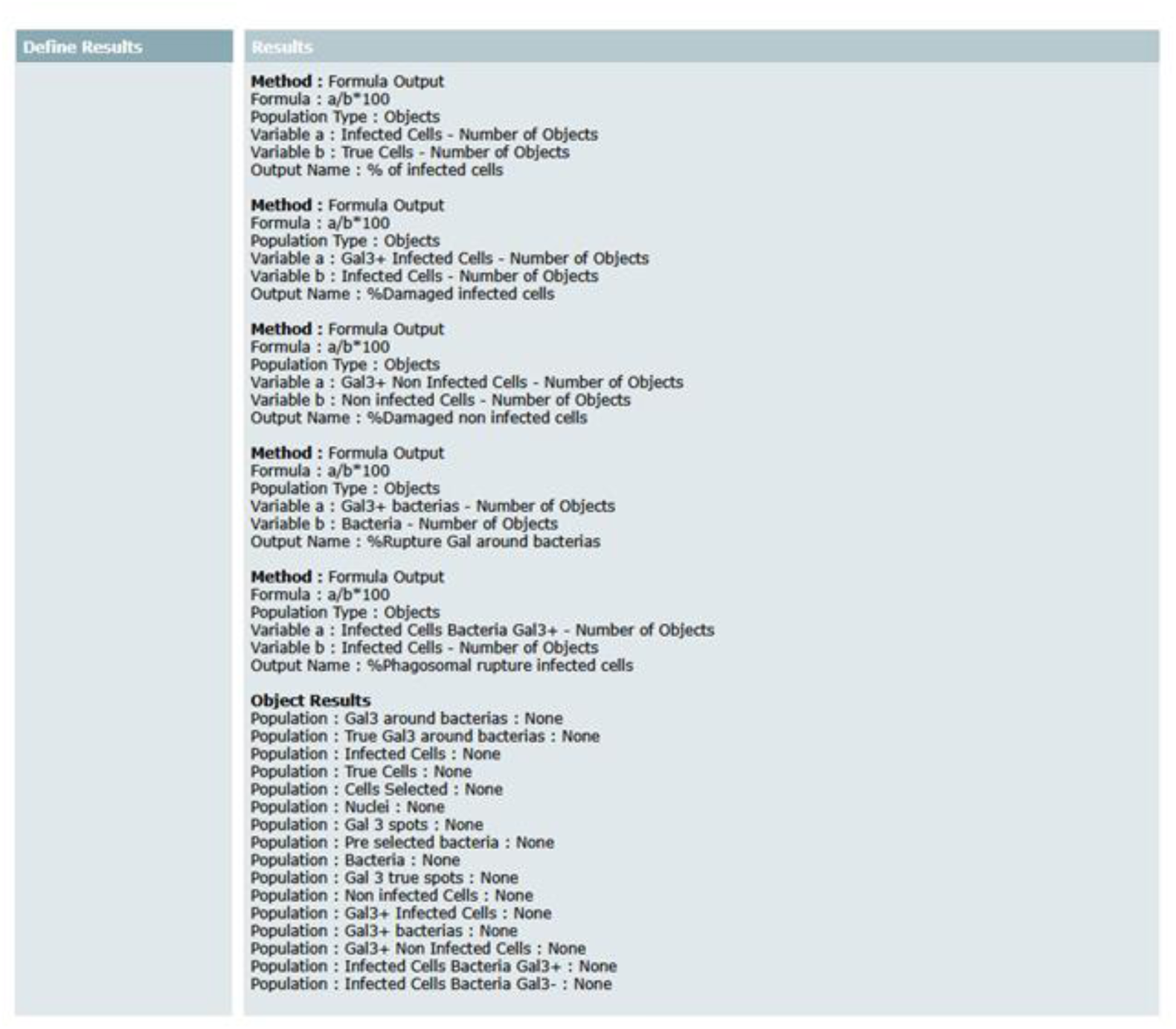
Image analysis workflow on Columbus software for the quantification of the colocalization of bacteria and Galectin-3.

